# Combined drought resistance strategies and the hydraulic limit in co-existing Mediterranean woody species

**DOI:** 10.1101/2022.04.01.486704

**Authors:** Asaf Alon, Shabtai Cohen, Regis Burlett, Uri Hochberg, Victor Lukyanov, Ido Rog, Tamir Klein, Herve Cochard, Sylvain Delzon, Rakefet David-Schwartz

## Abstract

- Woody species employ various strategies to cope with drought stress. We investigated similarities and differences in response to chronic drought to understand resistance strategies in co-occurring Mediterranean species.
- We studied five predominant Mediterranean species; *Quercus calliprinos, Pistacia palaestina, Pistacia lentiscus, Rhamnus lycioides*, and *Phillyrea latifolia* over two summers at three sites with different aridities. We measured key hydraulic and osmotic traits related to drought resistance, including resistance to embolism (Ψ_50_), carbon isotope signature (δ^13^C), pre-dawn (Ψ_PD_) and mid-day (Ψ_MD_) water potentials, and native (Ψ_s_) and full turgor (П_0_) osmotic potentials.
- Significant differences among species appeared in resistance to embolism. The species also showed differences in the water potential plastic response over the dry season. This interspecific variation increased at the end of the dry season and resulted in very narrow hydraulic safety margins (HSM). Consequently, predicted loss of hydraulic conductivity revealed species with significant native embolism. Two of the species also had seasonal changes in osmotic adjustment.
- Our detailed analysis indicates that co-existing Mediterranean woody species combine various drought resistance strategies to minimize mortality risk. However, all of them risk mortality as they approach their hydraulic limit near the dry margin of their distribution.

## Introduction

Drought is projected to increase in intensity and duration in many regions worldwide, including the Mediterranean (Spinoni *et al*., 2018; Xu *et al*., 2019) a hot spot for biodiversity (Myers *et al*., 2000). The current massive tree mortality in parts of the region is a major cause of concern for the extinction of native species (García de la Serrana *et al*., 2015; Cramer *et al*., 2018). Deciphering drought resistance strategies and their limitations in Mediterranean woody species is crucial for understanding changes in structure and function of plant communities threatened by climate change, and will help improve forests and woodlands’ sustainable management programs (Trumbore *et al*., 2015).

The various strategies used by woody species to cope with drought stress can be divided into three categories; escape, avoidance and tolerance (Delzon, 2015; Volaire, 2018). **Escape** is the temporary shedding of leaves and branches through which water is lost. **Avoidance** is actively minimizing water loss by stomatal closure or increasing water uptake through deep roots. **Tolerance** is maintaining physiological functionality during water loss, mainly by increased xylem resistance to embolism and osmotic adjustment to prevent turgor loss at the cell level. Woody species vary in their strategies to cope with drought, especially under natural conditions of prolonged and severe drought. Therefore, field studies of natural populations for a wide range of species and traits, and along aridity gradients, are necessary.

Drought can lead to embolism, a process that occurs via cavitation events in the xylem and causes hydraulic dysfunction by disrupting water conduction in the xylem (Tyree & Zimmermann, 1983). Embolism resistance is often expressed as the value of xylem water potential (Ψ_x_) corresponding to 50 or 88 percent loss of conductivity (PLC), Ψ_50_ and Ψ_88_, respectively (Tyree & Sperry, 1989). Embolism has been shown to be one of the leading causes of tree mortality worldwide (Anderegg *et al*., 2016; Adams *et al*., 2017). The variation in embolism resistance between species is large, and it appears that species habitat dryness plays a significant role in this variation (Maherali *et al*., 2004; Delzon *et al*., 2010; Choat *et al*., 2012; Larter *et al*., 2017; Skelton *et al*., 2018). As opposed to the large interspecific variation, it seems that intraspecific variation in resistance to embolism is limited. However, an analysis of 46 species suggests that significant intraspecific variation may occur (Anderegg, 2015).

Leaf water potential (Ψ_l_) is an indicator of plant water status. Pre-dawn leaf Ψ (Ψ_PD_) is measured when the plant is in equilibrium with soil water, and is a measure of the soil water availability as perceived by the plant. Ψ_PD_ is affected by drought severity and root depth (Nardini *et al*., 2016). The difference between either the Ψ_50_ or Ψ_88_ value and the minimum water potential observed in field conditions (Ψ_min_) is defined as the hydraulic safety margins (HSM) (Meinzer *et al*., 2009; Martin-StPaul *et al*., 2017). A narrow HSM means proximity to thresholds where there is a risk of hydraulic failure. A meta-analysis showed that most plants that resist drought do so by stomatal closure at a much higher water potential than that which causes hydraulic failure (Martin-StPaul *et al*., 2017). This suggests that species in nature avoid hydraulic failure by sacrificing photosynthesis (Meinzer *et al*., 2009; Johnson *et al*., 2011; Martin-StPaul *et al*., 2017; Creek *et al*., 2020).

Photosynthetic performance is usually measured in ecological studies through leaf carbon isotope discrimination, δ^13^C (Cernusak *et al*., 2013). Leaf δ^13^C is used to assess leaf gas exchange characteristics in C3 terrestrial plants. It is commonly used as a proxy for intrinsic water use efficiency (WUEi), which is the ratio between carbon assimilation (A) and stomatal conductance (g). WUEi is considered an integrative, long-term evaluation of photosynthetic performance rather than an instantaneous measurement (Dawson *et al*., 2002).

Stomatal closure has been shown to be correlated with the cell turgor-loss-point (TLP) (Mencuccini *et al*., 2015; Bartlett *et al*., 2016; Martin-StPaul *et al*., 2017). To keep turgid cells, plants may invest energy to reduce the cell TLP during dehydration. The water potential for TLP (Ψ_TLP_) is reduced through active accumulation of solutes, i.e. osmotic adjustment (Bartlett *et al*., 2012). The seasonal course leaf osmotic potential reveals the plasticity of osmotic adjustment as related to environmental changes (Bartlett *et al*., 2012).

Studies focusing on leaf traits of co-existing woody species in Mediterranean climates reveal variation in response to drought during the long rainless summer. Leaf defoliation in response to extreme drought was observed in *Juniperus phoenicea, Rosmarinus officinalis*, and *Rhamnus lycioides* but not in four other co-existing species (Gazol *et al*., 2017). Dehydration avoidance via stomatal closure at the cost of low carbon assimilation was demonstrated by *Pinus nigra*, while neighboring *Quercus ilex* and *Quercus faginea* showed dehydration tolerance via osmotic adjustment. The observed osmotic adjustment was more robust in the evergreen *Q. ilex* than in the semi-deciduous *Q. faginea*, explaining its better resistance to drought (Forner *et al*., 2018a). However, *Q. faginea* responded to intensified drought conditions by a more robust plastic stomatal response than *Q. ilex* (Klein *et al*., 2013; Forner *et al*., 2018b). Independent studies that measured δ^13^C agreed that pine exhibited better WUEi than co-existing oak species under prolonged drought or inter-annual precipitation differences. A significant variation in stomatal regulation, detected through δ^13^C and oxygen isotope composition in ten co-existing species in Spain, further emphasized the contrasting WUEi among species (Moreno-Gutiérrez *et al*., 2012). Combined analysis of Ψ_PD_ and sap flow of five species allowed designating *Pinus halepensis, Pistacia lentiscus*, and *Erica multiflora* as water savers, versus *Quercus coccifera* and *Stipa tenacissima* as water spenders (Chirino *et al*., 2011).

The above studies imply interspecific differences in the plastic stomatal response or osmotic adjustment; however, resistance to embolism, a key trait in species resistance to drought, was missing from those studies. Large variation in resistance to embolism in Mediterranean climates was found among 19 fynbos species in South Africa (Pratt *et al*., 2012) and nine chaparral species of the Rhamnaceae in California (Pratt *et al*., 2007) Both studies concluded that resistance to embolism is linked to the species’ post-fire recruitment strategy. In a study conducting year-round measurements of resistance to embolism, water potential, and Ψ_TLP_ in three co-existing species (Väänänen *et al*., 2019), it was found that *Phillyrea latifolia* tolerates drought via xylem resistance to embolism and osmotic adjustment, while co-existing *Pistacia lentiscus* and *Quercus calliprinos* avoid drought via stomatal regulation. However, these results did not explain the high mortality rate of *Q. calliprinos* compared to the other species, suggesting that an additional mechanism might occur in these co-existing species. This emphasizes the importance of evaluating a range of traits, species, and aridity gradients to represent better each species’ total traits repertoire used for drought resistance.

Here, we tested the hypothesis that co-existing Mediterranean species differ in drought resistance strategies, however, all can respond to drought intensification through plastic traits. To this end, we studied five co-existing species at three sites with different aridity, one of which is near the dry margin of the species distribution. We measured predawn and midday water potentials during the dry season and δ^13^C at the end of the dry season, resistance to embolism at the end of the wet season, and native and full-turgor osmotic potentials as related to midday water potential over the course of the dry season. The resulting data set allowed us to evaluate the response of co-existing species to prolonged drought.

## Materials and methods

### Sites and species

The steep climatic gradient in Israel is governed by Mediterranean weather patterns, characterized by long, dry summers, and changes gradually from mesic-Mediterranean to arid from north to south (Tielbörger *et al*., 2014). Three sites were chosen to represent Mesic-Mediterranean (MM), Mediterranean (M) and Semi-arid (SA) climate conditions along the natural rainfall gradient (Table 1). The three sites were undisturbed for the last 40 years, and thus represent ecologically equilibrated environments. Climate data was taken from stations of the Israel Meteorological Service (IMS, ims.gov.il); station Michmanim for the MM site, station Ramat Hanadiv for the M site, and station Netiv HaLamed-Heh for the SA site. All sites were characterized by a prolonged summer-fall rainless dry period, from the end of April to October. Aridity indexes (mean annual precipitation divided by potential evapotranspiration) for the SA, M, and MM sites are 0.27, 0.39, and 0.46, respectively. Annual precipitation (from September to August) in 2017-2018 was 337, 624, and 700 mm, and in 2018-2019 was 504, 651, and 1039 mm at the SA, M and MM sites, respectively. Temperature also differed along the climate gradient and average daily maximum temperature in the summer was 35°C, 32°C, and 29°C, respectively (Fig. **1a**). Maximum VPD in the summer ranged from 4 to 2.5 kPa from SA to M, respectively (Fig. **1b**). Soil types varied between the sites and the degree of clayey soil decreased from north to south. Five predominant native woody species were selected for the research (Table 2), which co-existed in 2,500 square meter plots at each of the three research sites. The SA site is close to the dry southern limit of the Mediterranean zone and the studied species (Supporting Information Fig. 1) (Danin & Plitmann, 1987).

**Table 1:** Three selected sites with main geographical, edaphic, and climatic characteristics.

| Code Site | *AI | Latitude | Longitude | Elevation (m) | MAP * (mm) | SMT *(°C) | WMT *(°C) | Soil type** |
| --- | --- | --- | --- | --- | --- | --- | --- | --- |
| MM | 0.46 | 32°54'N | 35°20'E | 430 | 700 | 30 | 10 | TR |
| M | 0.39 | 32°33'N | 34°56'E | 80 | 550 | 32 | 16 | TR and R |
| SA | 0.27 | 31°40'N | 34°56'E | 300 | 400 | 31 | 17 | Bright and Brown R |

\* AI, Aridity Index is the ratio of the annual precipitation and potential evapotranspiration. MAP, mean annual precipitation; SMT, Average maximum daily temperature in July; WMT, Average maximum daily temperature in January. \*Data for 1981–2000, provided by the Israel Meteorological Service. \*\* Data taken from GIS site of Israel Agriculture office (https://moag.maps.arcgis.com/), TR and R correspond to Terra Rossa and Rendzina, respectively.

**Fig. 1:**
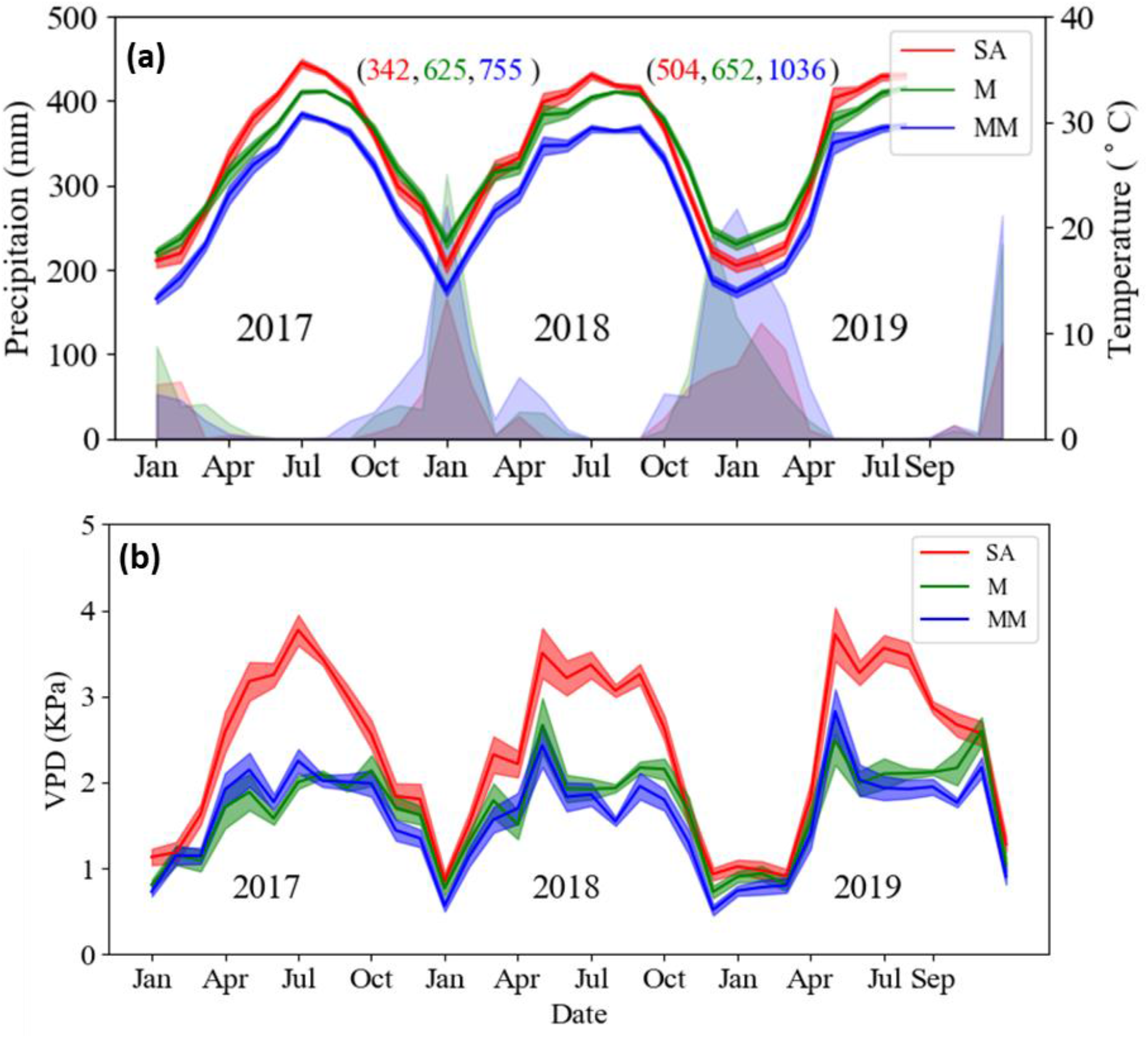
Meteorological data from the three study sites. (a) Monthly average daily maximum temperature and monthly precipitation. Numbers in brackets represent the annual precipitation for 2 consecutive winters (2017-2018, 2018-2019, calculated from September to September) for the SA (red), M (green), and MM (blue) sites, respectively. (b) Monthly average daily maximum vapor pressure deficit (VPD). Line shadow represents standard error.

**Table 2:**
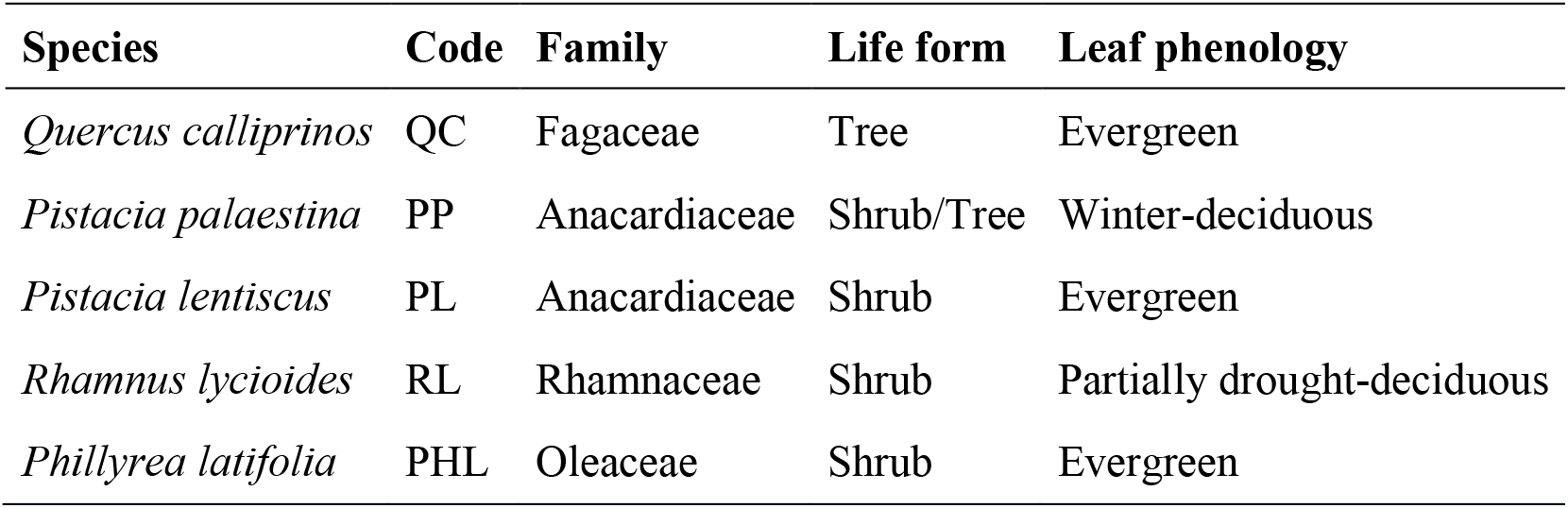
Characteristics of species selected for the research.

| Species | Code | Family | Life form | Leaf phenology |
| --- | --- | --- | --- | --- |
| *Quercus calliprinos* | QC | Fagaceae | Tree | Evergreen |
| *Pistacia palaestina* | PP | Anacardiaceae | Shrub/Tree | Winter-deciduous |
| *Pistacia lentiscus* | PL | Anacardiaceae | Shrub | Evergreen |
| *Rhamnus lycioides* | RL | Rhamnaceae | Shrub | Partially drought-deciduous |
| *Phillyrea latifolia* | PHL | Oleaceae | Shrub | Evergreen |

For all trait measurements described below, samples were taken from specific five labeled individuals per species at each site, unless otherwise stated.

### Hydraulic vulnerability measurements

Vulnerability curves (VC) for percent loss of conductivity (PLC) of stem samples as a function of water potential were measured in a Cavitron (Cochard, et al., 2005). This was done for all species except QC, which was not measured due to its long vessels (> 30 cm, personal data). It was impossible to find QC branches 1 m long for measurements in a nonstandard Cavitron with a large rotor diameter. Thus, we measured the PLC values of *Quercus coccifera*, which is an evergreen oak that belongs to the subgenus *Quercus* section *Cerris* and is considered a subspecies of QC (Toumi & Lumaret, 2010).

To avoid native embolism in branches that were tested for VC and PLC, samples were collected from the three sites at the end of the rainy season (April 2018), when water potentials were less negative than −1.5 MPa. Samples of RL at the MM site were not included in this analysis, as its branches were too short for the Cavitron. Two terminal branches (1 cm diameter and 100 cm in length) were harvested from the upper canopy of 5-7 individuals of each species and were sent in overnight mail to France (to Bordeaux and Clermont-Ferrand). VC curves were determined as previously described (Lamy et al., 2014). PLC was calculated every 1-2 MPa, following the equation:

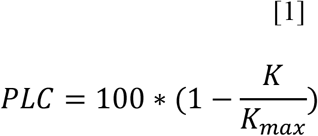

The sigmoidal curve was fitted to the following equation (Pammenter & Van der Willigen, 1998):

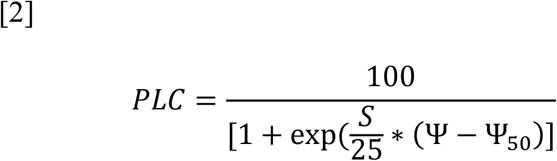

Where Ψ_50_ (MPa) is the xylem pressure inducing 50% loss of conductance and S (% MPa^-1^) is the slope of the vulnerability curve at the inflection point. Predicted PLC (PLC_P_) was calculated according to the actual leaf water potential and Equation 2.

### Field measurements of water potential

Field campaigns were conducted monthly at all sites during the rainless period (May - September) in two consecutive years; 2018, where predawn and midday water potentials were measured (Ψ_PD_ and Ψ_MD_, respectively) and in 2019, where only Ψ_MD_ was measured. The measurements were made in a Scholander-type pressure chamber (PMS, Corvallis, OR, USA). The decline in Ψ_MD_ in relation to soil dehydration (as reflected by Ψ_PD_) was analysed according to Meinzer et al. (2016). Regression lines were calculated for all species for each site. Slopes, Hydroscapes (which is a metric of stomatal control based on the area between the 1:1 line and the regression slope), and 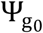 (extrapolated to find the value at which Ψ_PD_ = Ψ_MD)_ were calculated from the above analysis both for different sites for each species and for each species separately at all sites included (Supporting Information Table S8, S9). Ψ_min_ was taken as the lowest value of measured midday water potential in the field. Hydraulic safety margins (HSM) were calculated as Ψ_min_ - Ψ_x_, where Ψ_x_ is the xylem pressure inducing 12, 50 or 88% loss of branch hydraulic conductivity.

### Leaf δ^13^C

Carbon isotope ratio (δ^13^C) was measured in mature healthy sunlit leaves collected at the end of the dry period (August 2018) with a ^13^C cavity ring down analyzer (G2131i, Picarro, Santa Clara, CA, USA) as described by Nemera et al. (2020). Leaf intrinsic water-use efficiency (WUEi) was calculated using the species mean based on a leaf-scale model of C3 photosynthetic isotope discrimination (Farquhar et al., 1989):

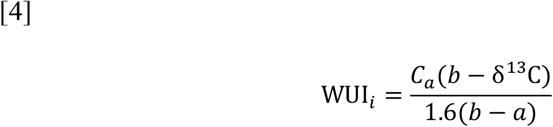

Where C_a_ is the atmospheric CO_2_ concentration in PPM and a and b are fractionation factors occurring during diffusion of CO_2_ through stomata pores (4.4‰) and enzymatic carbon fixation by Rubisco plus a small component accounting for mesophyll conductance (27‰), respectively.

### Osmotic potential

Leaf samples in which water potential was measured were frozen in liquid nitrogen for native osmotic potential (Ψ_s_) measurements. For full turgor osmotic potential (П_0_) measurements, an additional shoot, harvested from each individual, was cut under water, rehydrated for 2 hours, and measured in the pressure chamber to verify rehydration. For both cases (Ψ_s_ and П_0_), samples were packed into 250 µl tubes and were frozen in liquid nitrogen. Upon thawing, holes were drilled in the bottom of the frozen tubes, which were then put into other clean tubes that collected the liquid when centrifuged at 15000 RCF (g) for 2 min. Ten microliters from each Osmolality (mmol) of the samples was assessed with a vapor-pressure osmometer (VAPRO 5520 Wescor, Logan, UT). Conversion to pressure units was done by Van’t Hoff equation:

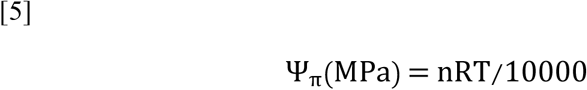

in which n is the solute concentration in mol/L, R is the universal gas constant (8.314472 L bar K^-1^ mol^-1^), and T is the temperature in °K. Temperature was taken as 25^0^; conversion ratio was 403.33 mmol/MPa.

Ψ_s_ and П_0_ were analyzed by correlation with the Ψ_MD_ values, and by covariance analyses which were performed to test the influence of Ψ_MD_, site, and the interaction between Ψ_MD_ and site on Ψ_s_ and П_0_. Slope regression analysis of Ψ_s_ Vs. Ψ_MD_ is a proxy for the osmotic potential due to both cell shrinkage and osmotic adjustment (OA, i.e., active solute accumulation). Slope regression analysis of П_0_ Vs. Ψ_MD_ is a proxy for OA only.

### Species characterization by drought-resistance strategies

Tolerance and Avoidance were quantified as numbers between 0 and 100 (less to most, respectively) for each species. Each strategy was evaluated from the below measured parameters which were converted to normalized values (NV) as follows:

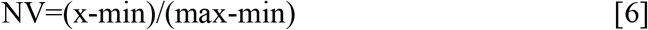

Where x is the measured value, and min and max are the minimum and maximum thresholds of each parameter.

“**Tolerance**” refers to “Xylem tolerance”, and was taken to be a normalized value of Ψ_50_. Values for normalization were from 0 to −18 MPa, the latter being the most negative Ψ_50_ reported (Larter et al., 2015).

“**Avoidance**” was calculated as the average of “Water access”, “Stringency of stomatal control”, and “Osmoregulation”. “Water access” was normalized from the minimum seasonal Ψ_PD_ at the SA site. Values for normalization were from 0 to −10 MPa. Normalized values were subtracted from 1. “Stringency of stomatal control” was normalized from Hydroscapes (HS), calculated from Ψ_PD_ vs. Ψ_MD_ according to Meinzer et al. (2016), where the full range of values was from 0 to 10 MPa. Normalized values were subtracted from 1. “Osmoregulation” was normalized from the slope of Ψ_MD_ vs. П_0_. Values for normalization were from 0 to 1.

“**Escape**” was the rank for “Drought-deciduous”. “Drought-deciduous” was evaluated from the literature (1, 0.5, and 0, refer to full-, partial-, and non-deciduous, respectively). Among the studied species, only RL is known to be partially drought-deciduous (Gazol et al., 2017).

### Statistical Analysis

Analysis of variance (ANOVA) was used (Python software, Python Software Foundation; JMP 14 software, SAS Institute Inc., Cary, NC, USA) to identify significant differences between species and sites. The Tukey-Kramer post hoc test was used to compare the results. Data fitting was carried out using Python software. Bartlett’s test for homogeneity of variances, using JMP 14 software was used to compare interspecific variation among sites and along the season within sites. Analysis of co-variance was used to test influence of both site and Ψ_MD_ on П_0_ and Ψ_s_, and also used to test influence Ψ_PD_ and site on Ψ_MD_.

## Results

### Effect of environmental drought on hydraulic traits

#### Resistance to embolism

Large differences in resistance to embolism were found between species, with the highest Ψ_50_ for PP (∼ −5 MPa) and lowest for PHL (< −10 MPa). All parameters of vulnerability curves for resistance to embolism per species, including slope, Ψ_12_, Ψ_50_, and Ψ_88_, were similar at the three sites (Fig. **2**, Table 3, Supporting Information Tables S1-S6), and were not influenced by the site, as shown by a two-factorial ANOVA (Table 3).

**Fig. 2:**
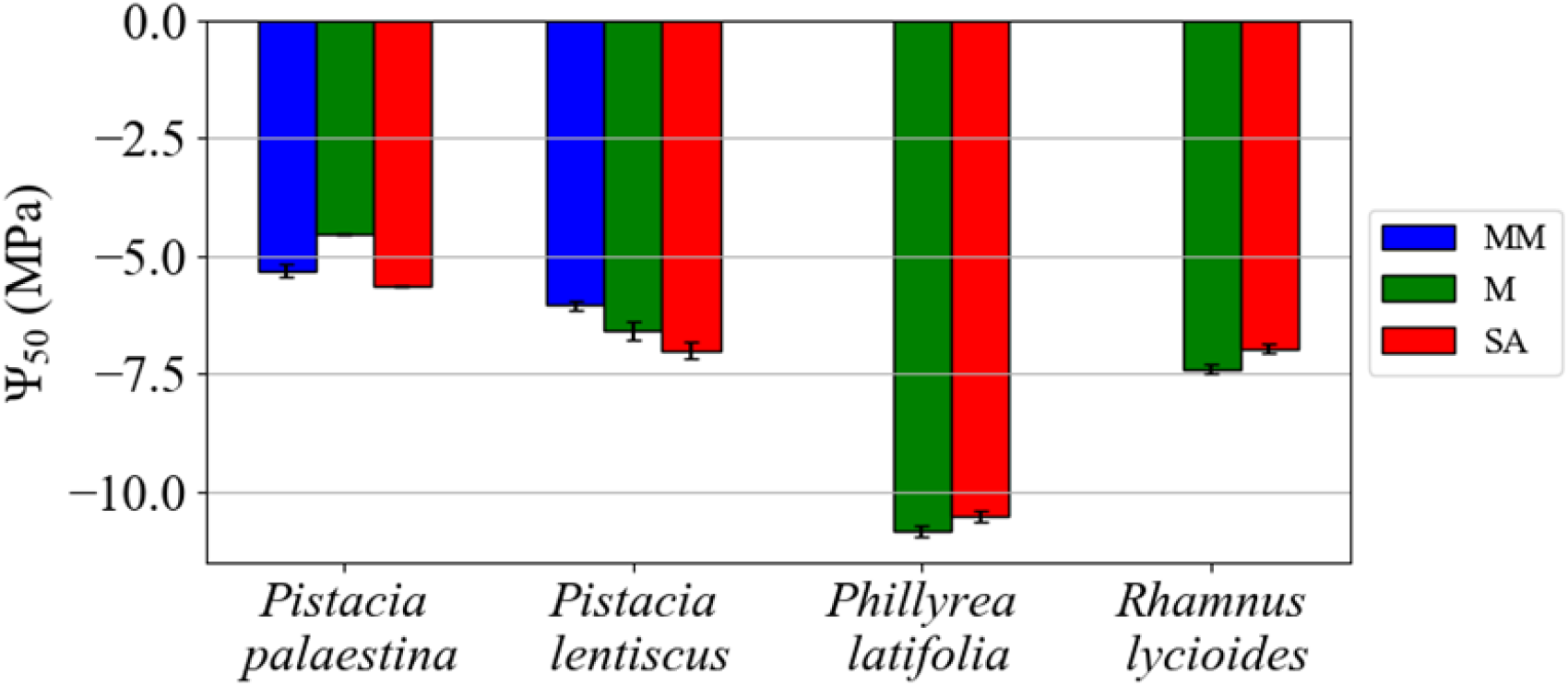
Ψ_50_ at the three sites (MM, Mesic-Mediterranean; M, Mediterranean; SA, Semi-arid). *Quercus calliprinos* was not measured due to excessive vessel length. Differences between species were significant, but no significant differences were observed between sites. Error bars indicate standard error (n = 5-7).

**Table 3:** Two-factorial ANOVA of the effect of species, site and the interaction between site and species on all measured traits. For all embolism resistance parameters the 2 factors analysis was done on four species (PL, PP, PHL, and RL) for two sites (M and SA). For the other parameters, analysis was done on four species (QC, PL, PP, and PHL) for three sites. Bold P values indicate statistically significant result.

| Trait | Species | Site | Species*Site |
| --- | --- | --- | --- |
| $\Psi_{50}$ | F(3,19)=84.5056<br>**,P=<0.0001** | F(1,19)=0.4501<br>,P=0.508 | F(3,19)=1.4025<br>,P=0.2636 |
| $\Psi_{12}$ | F(3,19)=17.4657<br>**,P=<0.0001** | F(1,19)=0.0105<br>,P=0.919 | F(3,19)=0.6733<br>,P=0.5759 |
| $\Psi_{88}$ | F(3,19)=27.4226<br>**,P=<0.0001** | F(1,19)=0.719<br>,P=0.4039 | F(3,19)=0.1714<br>,P=0.9148 |
| Slope | F(3,15)=0.6424<br>,P=0.5944 | F(1,15)=0.0095<br>,P=0.9229 | F(3,15)=0.9251<br>,P=0.442 |
| $\Psi_{min}$ (2018) | F(3,75)=84.7131<br>**,P=<0.0001** | F(2,75)=68.5311<br>**,P=<0.0001** | F(6,75)=4.9622<br>**,P=0.0003** |
| $\Psi_{min}$ (2019) | F(3,39)=92.7639<br>**,P=<0.0001** | F(2,39)=19.4777<br>**,P=<0.0001** | F(6,39)=2.0137<br>,P=0.087 |
| Leaf $\delta^{13}C$ / WUEi | F(3,33)=2.4531<br>,P=0.0806 | F(2,33)=22.3469<br>**,P=<0.0001** | F(6,33)=1.3517<br>,P=0.2629 |
| PLCp (2018) | F(3,75)=61.1677<br>**,P=<0.0001** | F(2,75)=13.0761<br>**,P=<0.0001** | F(6,75)=8.4556<br>**,P=<0.0001** |
| PLCp (2019) | F(3,39)=62.9183<br>**,P=<0.0001** | F(2,39)=9.6769<br>**,P=0.0004** | F(6,39)=6.3173<br>**,P=0.0001** |
| HSM<sub>12</sub> (2018) | F(3,75)=109.954<br>**,P=<0.0001** | F(2,75)=57.7828<br>**,P=<0.0001** | F(6,75)=10.5175<br>**,P=<0.0001** |
| HSM<sub>12</sub> (2019) | F(3,75)=110.3402<br>**,P=<0.0001** | F(2,75)=48.928<br>**,P=<0.0001** | F(6,75)=10.6456<br>**,P=<0.0001** |
| HSM<sub>50</sub> (2018) | F(3,75)=111.7792<br>**,P=<0.0001** | F(2,75)=40.8695<br>**,P=<0.0001** | F(6,75)=11.3523<br>**,P=<0.0001** |
| HSM<sub>50</sub> (2019) | F(3,39)=87.4207<br>**,P=<0.0001** | F(2,39)=17.8401<br>**,P=<0.0001** | F(6,39)=5.0945<br>**,P=0.0006** |
| HSM<sub>88</sub> (2018) | F(3,39)=88.1662<br>**,P=<0.0001** | F(2,39)=14.9253<br>**,P=<0.0001** | F(6,39)=5.0271<br>**,P=0.0007** |
| HSM<sub>88</sub> (2019) | F(3,39)=89.8414<br>**,P=<0.0001** | F(2,39)=12.7135<br>**,P=<0.0001** | F(6,39)=5.5257<br>**,P=0.0003** |
| $\Psi_{PD}$(Sep_2018) | F(3,74)=73.7787<br>**,P=<0.0001** | F(2,74)=39.6804<br>**,P=<0.0001** | F(6,74)=2.3523<br>**,P=0.0391** |
| $\Psi_{PD}$(May_2018) | F(3,32)=17.1726<br>**,P=<0.0001** | F(2,32)=85.8451<br>**,P=<0.0001** | F(6,32)=21.302<br>**,P=<0.0001** |
| $\Psi_s$ (June 2019) | F(3,11)=10.7141<br>**,P=<0.0001** | F(2,11)=0.3355<br>,P=0.7179 | F(6,11)=1.8519<br>,P=0.1263 |
| $\Pi_0$ (June 2019) | F(3,11)=21.5185<br>**,P=<0.0001** | F(2,11)=7.5033<br>**,P=0.0033** | F(6,11)=4.9563<br>**,P=0.0024** |

#### Leaf water potential

Minimum Ψ_MD_ was found at the end of the season at the SA site. For QC and PHL minimum values were close to, but did not decline below Ψ_12_, while for the other species minimum Ψ_MD_ values were significantly lower than Ψ_12_ (Fig. **3**). Significant differences in Ψ_PD_ between sites for each of the species were found in each sampling along the dry season (Fig. **3**, Supporting Information Table S7). A strong influence of site on Ψ_PD_ was found at the beginning and end of the dry season (Table 3). Ψ_PD_ was more negative at the SA than at the MM and M sites (Fig. **3a-e**, Supporting Information Table S7). Ψ_min_ was significantly influenced by site in both 2018 and 2019 (Table **3**, Fig. **3f-g**).

**Fig. 3:**
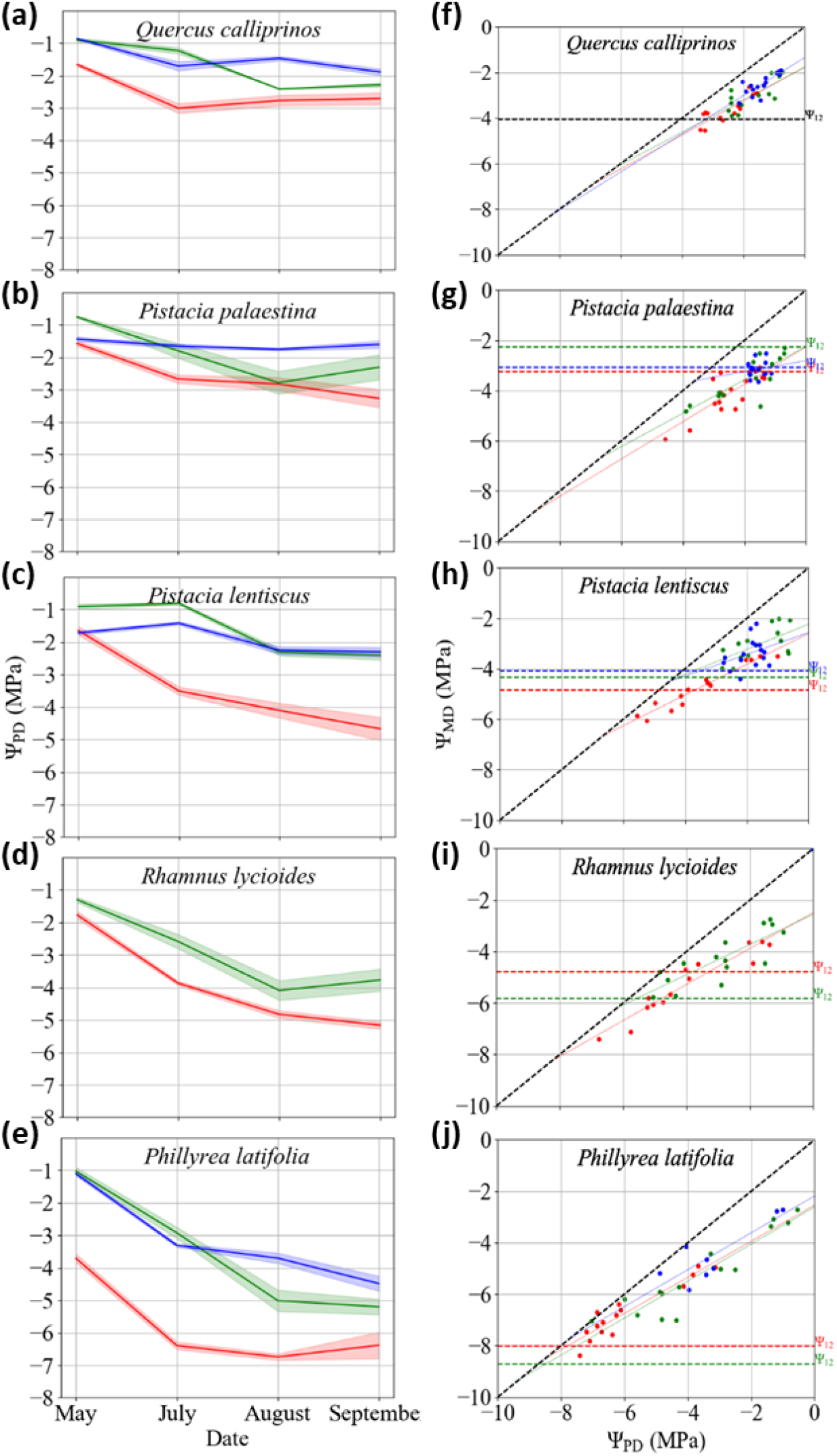
Leaf water potential values during the dry season of 2018 at the Mesic-Mediterranean (blue), Mediterranean (green), and Semi-arid (red) study sites. **a-e**; Courses of predawn leaf water potential for each species. Line shadow represents standard error. **f-j**, Midday water potential (Ψ_MD_) vs. predawn water potential (Ψ_PD_) for all species during the dry season. Each point on a plot is the average of two twigs. Regression parameters can be found in Table S3. Thick dashed black line represents 1:1 ratio. The thin colored lines are the regressions of all points of each site. Vertical dashed lines indicate the point of incipient embolism (Ψ_12_) values for the MM (blue), M (green), and SA (red) sites. Vertical dashed black line in **f** indicate point of incipient embolism based on *Q. coccifera* data (see Material and Methods section).

#### HSM and PLCp

The HSM in 2018 was narrower at the SA site than at the M and MM sites for three of the species, while the other two species had narrower HSM’s at the SA and M site in comparison to the MM site (Fig. **4a,b**, Supporting Information Tables S1, S3, S5). These differences were less prominent in 2019 (Fig. **4d,e**, Supporting Information Tables S2, S4, S6). Values of HSM_50_ less than 1 MPa were found at the SA site in 2018 (for PP and RL, Supporting Information Table S1), and the predicted PLC (PLCp) for those species reached values of 30% or more (Fig. **4**). In both 2018 and 2019 HSM_12_ declined to negative values at the SA site (Fig. **4a,e**, Supporting Information Tables S3, S4). The calculated HSM of Ψ_12_, Ψ_50_, and PLCp were affected by site in 2018, but not in 2019, which was a wetter year (Table 3, Fig. **1**).

**Fig. 4:**
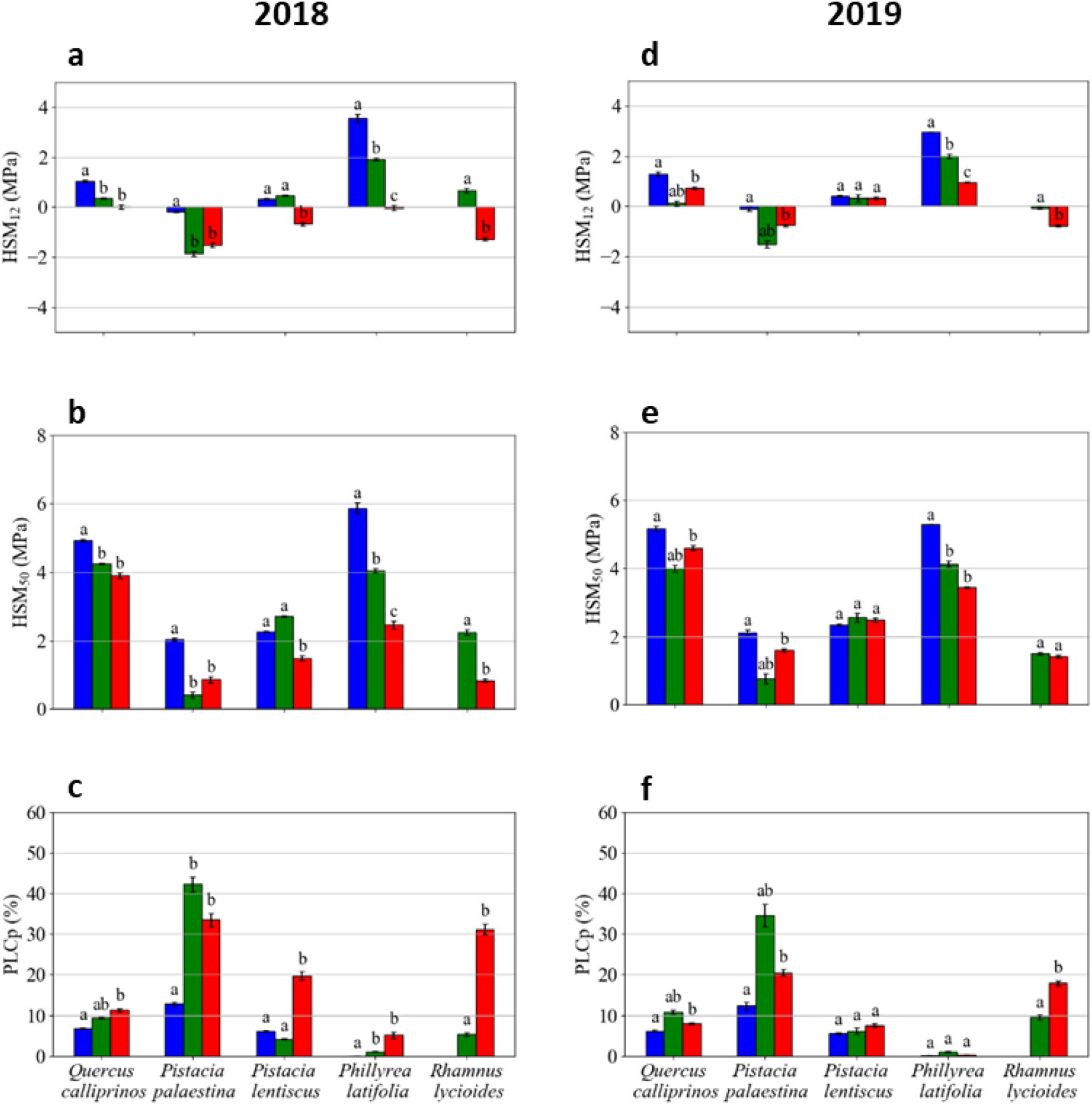
HSM- Hydraulic safety margins (HSM12 and HSM50) and PLCp (% of predicted embolism) for all species at the different sites, a-c represent 2018 data, d-f represent 2019 data. MM, Mesic-Mediterranean; M, Mediterranean; SA, Semi-arid. Different lowercase letters denote significant differences among sites. HSMs of QC are based on the PLC curve of *Q. coccifera* (See Material and Methods).

#### Carbon-water balance

Leaf δ^13^C and its derivative WUEi was higher at the drier site than at the wetter sites. Differences were significant for two of the species (Fig. **5**). The site influence on leaf δ^13^C was significant in the two-factorial ANOVA (Table 3).

**Fig. 5:**
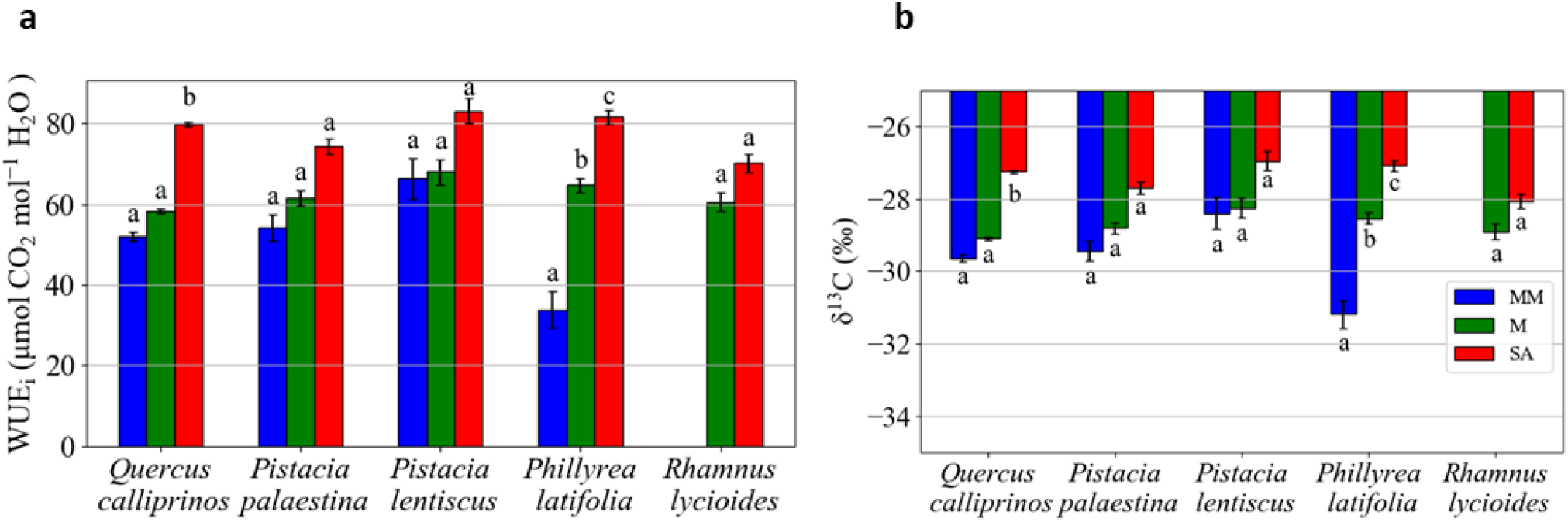
Leaf δ^13^C (a) and the derived WUEi (b) of the leaves of five species at the study sites at the end of the dry season. MM, Mesic-Mediterranean; M, Mediterranean; SA, Semi-arid. Different lowercase letters denote significant differences among sites.

#### Osmotic potential

Changes in osmotic potential were significant for all the species along the dry season, in relation to the decline in Ψ_MD_ (Fig. 6 f-j, Table S14). Covariance analysis for Ψ_s_ revealed a significant site effect only for PL, and a significant Ψ_MD_ effect for all species (Supporting Information Table S14). Osmotic adjustment differed substantially among species, being large in PHL and QC and minor in RL (which was expressed in significant Ψ_MD_ effect) negligible in PL, and nonexistent in PP (Fig. **6f-j**, Supporting Information Table S10). Covariance analysis for П_0_ revealed a significant site effect only for PL and PHL (Supporting Information Table S15).

**Fig. 6:**
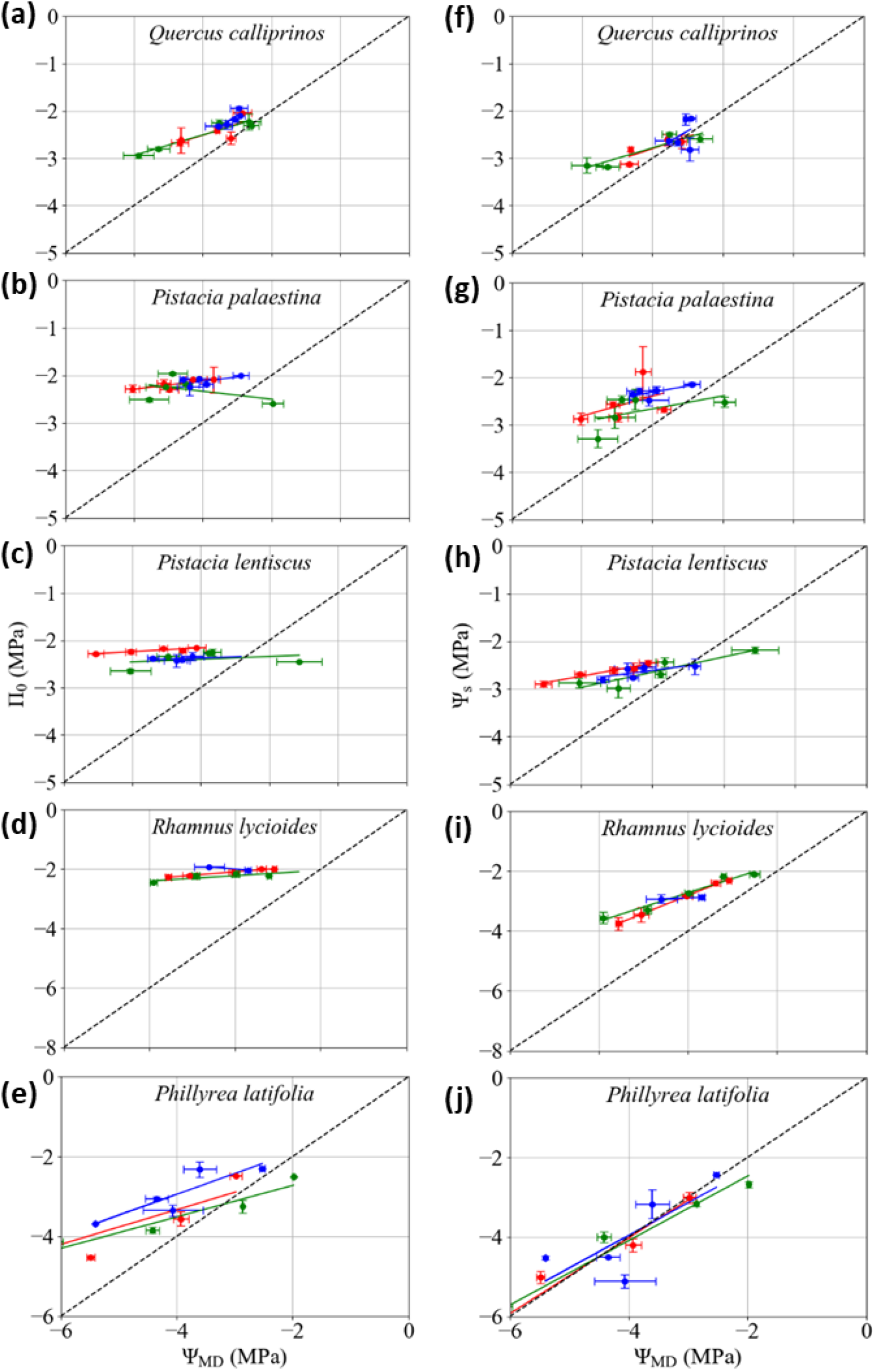
Osmotic potential and leaf water potential for all species measured in summer 2019 at the Mesic-Mediterranean (blue), Mediterranean (green), and Semi-arid (red) sites. **a-e**: Leaf water potential at midday (Ψ_MD_) vs. osmotic potential at full turgor (П_0_). f-j, Leaf water potential at midday (Ψ_MD_) vs. native osmotic potential (Ψ_s_). Regression parameters can be found in Supporting Information Table S10.

Accordingly, most of the reduction in Ψ_s_ during the dry season in QC is from active osmolyte accumulation, while in PL, PP and RL most of the reduction is due to cell shrinkage. PHL seems to retain both mechanisms.

### Species comparison by trait

#### Resistance to embolism

Resistance to embolism, expressed by Ψ_50_ (Fig. **2**), Ψ_12_, and Ψ_88_, showed interspecific variation at the SA and M sites (Table 3, Supporting Information Tables S1-S6). PHL demonstrated the most negative values followed by RL and PL, while PP showed the least negative value.

#### Water potential

Interspecific variations were evidenced at each site, and at each measurement event (Supporting Information Tables S11, S12). PHL had the most negative Ψ_PD_, followed by RL and PL, while QC and PP had the highest Ψ_PD_ values (Fig. **3a-e**). Two-factorial ANOVA showed that Ψ_min_, which is the lowest Ψ_MD_ value at the end of the dry season, was affected by the species and site. However, it was affected by the interaction of species-on-site only in 2018 (Table 3).

A comparison of the interspecific variation among sites using Bartlett’s test for homogeneity of variances, resulted in a significant difference (*P* = 0.0259) between sites at the beginning of the dry season. Interspecific variation was evidenced at the SA site more than at the M and MM sites (Table 4). The rest of the sampling dates did not show a significant difference in interspecific variation between sites; however, the SA site had higher values than the M and MM sites (Table 4).

**Table 4.**
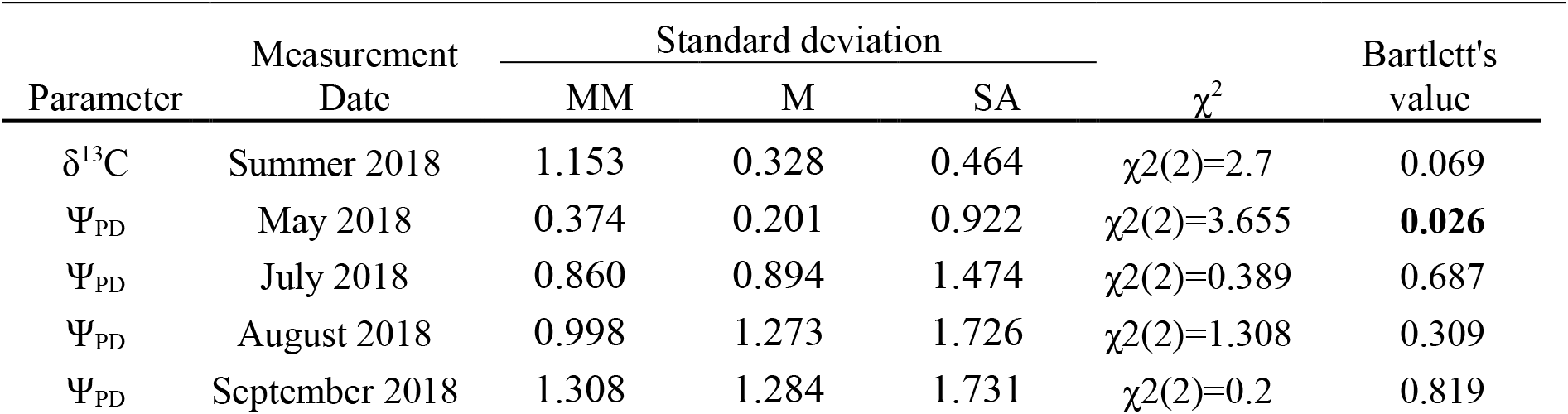
Bartlett’s test for homogeneity of variances for all species per date at the different sites. Bold indicates significant difference.

| Parameter | Measurement Date | Standard deviation | | | $\chi^2$ | Bartlett's value |
| --- | --- | --- | --- | --- | --- | --- |
|  |  | MM | M | SA |  |  |
| $\delta^{13}\text{C}$ | Summer 2018 | 1.153 | 0.328 | 0.464 | $\chi^2(2)=2.7$ | 0.069 |
| $\Psi_{\text{PD}}$ | May 2018 | 0.374 | 0.201 | 0.922 | $\chi^2(2)=3.655$ | **0.026** |
| $\Psi_{\text{PD}}$ | July 2018 | 0.860 | 0.894 | 1.474 | $\chi^2(2)=0.389$ | 0.687 |
| $\Psi_{\text{PD}}$ | August 2018 | 0.998 | 1.273 | 1.726 | $\chi^2(2)=1.308$ | 0.309 |
| $\Psi_{\text{PD}}$ | September 2018 | 1.308 | 1.284 | 1.731 | $\chi^2(2)=0.2$ | 0.819 |

Species showed different evolution of Ψ_PD_ along the dry season, and reached different values at the end of the season (Fig. **3a-e**). In each site, the interspecific variation increased during the dry season (Table 4), however, a comparison of the variations along the season at the different sites (using Bartlett’s test) found a significant (*P* = 0.0258) increase in interspecific variation only at the M site (Table 5). Slopes of Ψ_PD_ along the dry season were different between species in each site, while the higher value was that of RL and PHL, followed by PL, PP, and QC (Supporting Information Table S18).

**Table 5.**
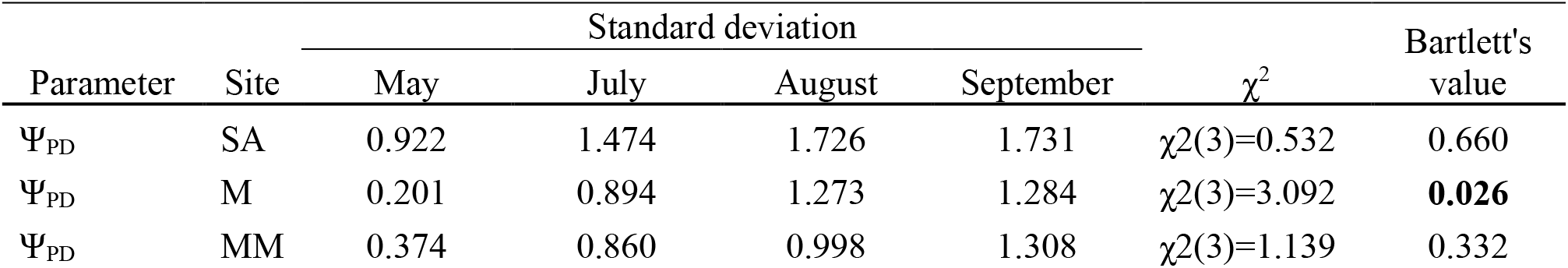
Bartlett’s test for homogeneity of variances for all species per site at the different dates. Bold indicates significant difference.

#### HSM

Similar to Ψ_PD_, the HSM of Ψ_50_, Ψ_12_, and Ψ_88_ showed interspecific variation (Tables 3, Supporting Information Tables S1-S6). PP (at SA and M sites) and RL (at SA site) had a very narrow HSM, i.e. less than 1MPa (Fig. **4**). However, in 2019, which was a wetter year (Fig. **1**), the HSM of all species was wider than in 2018. In 2018, the HSM based on Ψ_12_ at the SA site reached negative values for all species, except for QC, which had 0.02±0.21 MPa (Fig. **4a**, Supporting Information Table S3). In 2019, negative HSM values for Ψ_12_ were recorded only for PP and RL at all sites (Fig. **4d**, Supporting Information Table S4).

#### Carbon-water balance

One-way ANOVA suggested differences between species in leaf δ^13^C and WUEi per site (*P* = 0.0682, Table 3, Supporting Information Table S13). A comparison of the interspecific variation among sites using the Bartlett’s test suggested a strong tendency to significance (*P* = 0.0693), with the MM site showing the highest interspecific variation, while SA and M sites showed reduced variation (Table 4). These results were attributed to PHL and QC, which had significant differences in leaf δ^13^C between sites (Fig. **5a**).

#### Osmotic potential

Osmotic potential differed significantly between species in Ψ_s_ and П_0_ (Table 3). In addition, significant differences between species were revealed by covariance analysis for the different species at different sites (Supporting Information Tables S16, S17).

### Correlation between traits

#### The relationships between Ψ_PD_ and Ψ_MD_

Analysis of the various sites for each species showed that the slopes at the drier sites (M and SA) were steeper than the slope at the M site, except for QC. However, covariance analysis did not show significant differences between the slopes, except for PL. Hydroscape values for all species were always larger at the SA site than at the two wetter sites (Fig. **3f-j**, Supporting Information Table S8, S19). Analysis which includes the data from the various sites for each species showed that slopes ranged from 0.83 for QC to 0.68 for RL (Supporting Information Table S9). For QC and PP, the extrapolated values were several MPa lower than the lowest data points, −8.9 and −8.7 MPa, respectively.

However, for PHL, RL and PL the lowest points were close to the regression values at equality, −8.7, −7.6, and −7.1 MPa, respectively (Supporting Information Table S9). Hydroscape values were 10.7, 9.15, 7.31, 6.31, and 3.52 MPa^2^, for RL, PHL, PL, PP, and QC, respectively (Table S9).

### The relationships between osmotic parameters and Ψ_MD_

For all species significant linear correlations were found between osmotic potential, Ψ_s_, and Ψ_MD_ along the season (Fig. **6a-e**). The effect of Ψ_MD_ on Ψ_s_ was evident for all species, while a site effect was found only for PL (Supporting Information Table S14). A linear correlation was also found between П_0_ and Ψ_MD_ (Fig. **6f-j**, Supporting Information Table S10), while only QC, PHL, and RL had a Ψ_MD_ effect on П_0_ (Supporting Information Tables S15). There was evidence for a site effect on П_0_ for PHL and PL, while no effect for the interaction of Ψ_MD_ with site was revealed for any of the species (Supporting Information Table S15). The largest osmotic adjustment was observed for PHL, and ranged from −2 to −5 MPa along the dry season, with similar responses at all sites (Fig. **6j**). Osmotic adjustment for QC ranged from −2 to −3 MPa during the dry season, mostly at the M and SA sites (Fig. **6f**). A weak, but significant, osmotic adjustment was observed for PL and RL at the SA site (Fig. **6g-i**, Supporting Information Table S10).

### Species characterization by drought-resistance strategies

The results described above allowed the assessment of drought resistance strategies for each of the studied species (Fig. **7**). That was achieved by quantifying tolerance, avoidance, and escape strategies, as described in the Material and Methods (Supporting Information Table S19). Thus, we found that extreme resistance to embolism confers tolerance in PHL, which also uses osmotic adjustment to resist drought. Osmotic adjustment is a major trait in QC, which probably has deep roots to access water, supporting its stomatal opening during drought. PL and RL do not use osmotic regulation but have intermediate values of resistance to embolism and almost complete stomatal closure. PP has low resistance to embolism and no osmotic regulation, but its continued water uptake along the dry season suggests it has deep roots. RL is known to escape drought by partial leaf defoliation (Gazol et al., 20017).

## Discussion

The resulting data set of the current study, in addition to known species-specific characteristics, demonstrated that co-existing Mediterranean species minimize mortality risk by combining drought resistance strategies. The results support the hypothesis that regulation of water potential is a result of a robust plastic response, while resistance to embolism is a fixed rigid trait that is not affected by site aridity. These characteristics led to a decline in the HSM with increasing drought intensity leading to negative HSM in several cases and suggesting hydraulic failure at the dry SA site.

### Relating environmental factors to phenotype

The large data set provided by this study, encompassed three aspects of environmental drought, including site aridity, seasonal drought, and inter-annual climate differences. While all site characteristics were less favorable at the SA site than at the two wetter sites, precipitation and VPD differed the most, and actually played major roles in determining the drought intensity of the site. The rainless summer, together with the inter-annual precipitation differences, further emphasized the stress intensity that the studied species confronted. The SA site differed significantly from the other two sites for most of the traits, suggesting that species were closer to their physiological limits at the dry edge. The latter is in agreement with Feng, et al. (2019) and Guo, et al. (2020) who emphasized that temporal and spatial variability in the environment is important in determining plant response to drought, as opposed to characterization of species without considering environmental influences.

Our results show interspecific variation in resistance to embolism (Fig. **2**). However, no intraspecific variation was evident for this trait. A lack of intraspecific variation is in agreement with previous studies that showed similar resistance to embolism in distantly separated populations (Martínez-Vilalta et al., 2009; Lamy et al., 2014; González-Muñoz et al., 2018; Lobo et al., 2018; Li et al., 2019; Bittencourt et al., 2020). However, although the three tested sites differ in climate characteristics, the lack of intraspecific variation might also be due to the continuous geographical distribution of the tested species (Figure S1), which prevented differentiation between populations due to continuous gene flow. In addition, it is possible that other remote populations, which were not included in the current study, do possess intraspecific variation in resistance to embolism. Several studies have reported intraspecific variation in resistance to embolism in angiosperms. Examples are *Cordia alliodora, Artemisia tridentate, Fagus sylvatica, Populus trichocarpa*, and in Mediterranean and chaparral shrubs (Kolb & Sperry, 1999; Sparks & Black, 1999; Choat et al., 2007; Wortemann et al., 2011; Pratt *et al*., 2012; Jacobsen *et al*., 2014; Stojnić et al., 2017).

As opposed to the stability of resistance to embolism, the stomatal response was very plastic as reflected in changes in leaf water potential (Ψ_PD_ and Ψ_MD_) in response to drought in all species in relation to site aridity, seasonality, and inter-annual climate differences (Fig. **3**, Table 3). The difference between species in the Ψ_PD_ slope along the season, especially at the SA site (Fig.**3**, Table S18), suggests species differentiation in the degree of plasticity, where RL and PHL showed the strongest, and QC showed the least plastic response. The interspecific variation in Ψ_PD_ seemed to increase with drought and along the dry season, i.e., it was inversely related to aridity (Tables 4, 5).

The hydroscapes (HS, Supporting Information Tables S8, S9), which showed differences between species, can be used as proxies for stringency of stomatal regulation (Meinzer et al., 2016). They gave a nearly mirror image of plasticity, where QC had the most stringency of stomatal regulation, while PHL and RL had the least stringency (Fig. **3**, Supporting Information Tables S8, S9).

Variations in Ψ_PD_ and its plasticity between species along the dry season may reflect differences in root depth, where Ψ_PD_ of more deeply rooted species, such as QC, appears high with weak plasticity along the season (Crombie et al., 1988). Roots of five meter depth have been reported for PP (Jakoby et al., 2020), and QC (Canadell et al., 1996). Co-existing species with different root depths sustain niche segregation to share soil water resources (Palacio *et al*., 2017). Niche segregation has been shown in Mediterranean-type ecosystems through the leaf life span and Ψ_min_ as “anchor traits” among different morphological traits used to distinguish contrasting strategies of drought tolerance vs. avoidance(Ackerly, 2004). Our study, which focused on embolism resistance and leaf water potential, also suggests niche segregation in Mediterranean species. Another approach to exploring niche segregation divides species by different water use patterns (Redtfeldt & Davis, 1996), which has been recently shown to be important in increasing forest productivity and the carbon sink in semi-arid regions (Rog et al. 2021). Our study suggests that niche segregation is sustained under different drought conditions.

### The dearth of hydraulic safety margins near the dry edge of species distribution

The value of Ψ_L_ for complete stomatal closure 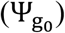 for QC and PP was more than 1 MPa lower than Ψ_min_, while for PL, RL and PHL Ψ_min_ was close to 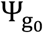 (Fig. **3f-j**). The proximity of the minimum Ψ_PD_ to 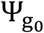 at the SA site indicates that these plants approached the point of null activity, which corresponds with the large reduction in Ψ_PD_ across sites, as shown in Fig. **3**.

The Ψ_50_ HSM seemed to change according to annual precipitation (MAP, Fig. **4**), especially at the SA site, suggesting that a drought year would have more impact on species vulnerability. This is in agreement with Ziegler et al. (2019), who showed that for tropical trees the Ψ_50_ HSM became narrower, but still positive, in dry years. In the current study, all species (except for QC) crossed the Ψ_12_ threshold (Fig. **3f-j**) and some degree of difference between Ψ_PD_ and Ψ_MD_ remained, suggesting that species maintain some stomatal opening when embolism is low. Interestingly, PP had a negative Ψ_12_ HSM in all sites in both years (Fig. **4**), emphasizing the trade-off between hydraulic safety and carbon assimilation in this winter-deciduous species. In addition, results of predicted PLC (PLCp, Fig. **4c,f**) suggest that all species experienced embolism, which increased with drought intensity. These results suggest that species approach the limit of hydraulic capacity at the site near the dry margins of their distribution.

Evidence for embolism of stem xylem in nature is rare. Johnson et al. (2018) recently reported measurements implying that negative HSM’s in *Quercus fusiformis* and *Prosopis glandulosa* occurred during the most severe drought in recorded history in central Texas. Fontes et al. (2018) found negative HSMs in *Eschweilera cyathiformis* and *Pouteria anomala* that experienced extreme drought during the strong El Nin∼ o that occurred across Amazonia in 2015–2016. The two above reports resulted from extreme climate events, while the species in our study seem to confront severe drought every year. Recent studies on *Prunus ramonensis* and *Pyrus syriaca* also reported potential embolism in nature (Paudel et al., 2019a; Paudel et al., 2019b). Taking a modeling approach, Benito et al. (2018) use minimum soil water potential data and HSMs of 44 European woody species and found that negative HSMs explain the mortality of 15 species at the driest margins of their distribution.

### Interspecific variation in response to drought

The interspecific variation in Ψ_PD_ increased with aridity and along the dry season (Tables 4, 5). This difference was supported by the significant effect of the species-by-site interaction on Ψ_PD_ (Table 3). However, this effect appeared only at the beginning of the dry season, where species operate at their relatively maximal physiological activity. As drought progressed, species reached minimum water potential, at which time the effect of site on differences was not significant.

The Ψ_PD_ results were supported by the leaf δ^13^C values that increased with site aridity, suggesting an increase in WUE_i_ (Fig. **5**) (Farquhar et al., 1989). Similar results were obtained by Rumman et al. (2018), who found a negative correlation between MAP and WUE_i_ for precipitation up to 1000 mm/year, above which the trend flattened, indicating that isotopic discrimination in wet environments remained nearly constant.

The current study shows that the interspecific variation in leaf δ^13^C tends to be larger at the MM site than at the M and SA sites (Table 4). This result suggests that genetic variation in carbon assimilation rate is more pronounced in environmental conditions favoring high stomatal conductance, as compared to the M and SA sites. It also suggests that species under severe environmental drought, that demonstrate different plasticity, reach similar minimum rates of carbon assimilation (Fig. **5**). This is in agreement with Forner et al. (2018b), who showed that the interspecific variation in leaf δ^13^C in three woody Mediterranean species in two consecutive wet years was reduced after an extremely dry year. Together, these results encourage the measurement of interspecific variation in carbon assimilation rates in wet rather than dry environments. Furthermore, high versus low interspecific variation in leaf δ^13^C could be a proxy for evaluating water stress in multi-species ecological niches.

As all the species in our study suffered from severe drought at the SA site, as manifested in a significant reduction in HSM’s, not all showed osmotic adjustment, suggesting that this mechanism is species dependent. Osmotic adjustment in drought has been reported for PHL (Serrano *et al*., 2005), and is also well known in Olive species, which are related to PHL (*Oleaceae* family) (Lo Gullo & Salleo, 1988; Sofo *et al*., 2008). It has also been found in *Quercus* species (Delıgö & Bayar, 2018; Aranda *et al*., 2020) but not in QC (as far as we know), and in PL (Álvarez *et al*., 2018) only in response to salinity.

The low degree of osmotic adjustment in RL and nonexistent osmotic adjustment in PP may also be related to their deciduous nature, and may result from a strategy of allocating fewer resources to the leaf, similar to the findings of Liu *et al*. (2011) who showed higher capacities of osmotic adjustment in evergreen shurbs than in decidous.

## Conclusions

As illustrated in Figure 7, each of the co-occurring species in our study combines drought-resistance strategies to minimize the risk of mortality. However, all approached the limit of their hydraulic capacity at the site near the dry margins of their distribution. The hydraulic limit was more pronounced in the drier year, suggesting that a slight reduction in precipitation is more likely to put species at the dry margins of their distribution at the risk of mortality.

**Fig. 7:**
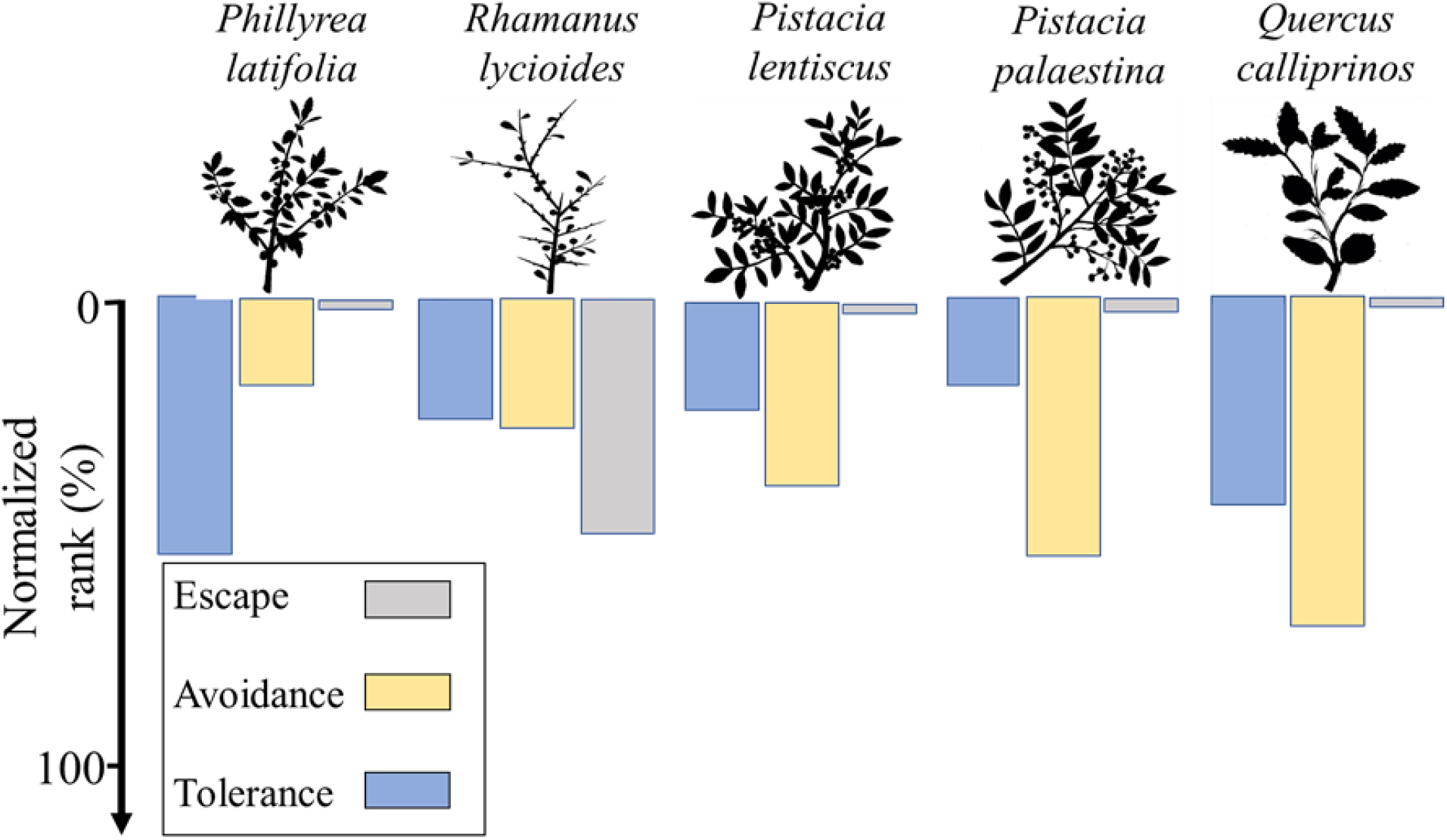
A superposition for the three different drought-resistance strategies (Tolerance, Avoidance, and Escape) for a species, evaluated as normal parameters (scaled from 0 to 100%) derived from measured parameters. Detailed evaluations are presented in Supporting Information Table S19. Drawing by Ilana Stein.

## Acknowledgments

We acknowledge the Ramat Hanadiv team for their administrative expertise and technical assistance. We thank Rotem Attias, Shai Tamari, Feng-Feng and Junzhou Liu for their help in the fieldwork. We thank Gaëlle Capdeville for assisting in the Cavitron measurements. We thank Hillary Voet for her statistical advice. We thank the three anonymous referees for providing useful comments, which improved the article. This work was supported by the Ministère des Affaires Etrangères et du Développement International (France) and the Ministry of Science (Israel), under the Research Program “Maïmonide-Israel” to RDS, SD, SC and HC.

## Author Contribution

AA, RDS, SC, SD, and HC designed experiments and interpreted data. AA, RB, and VL collected and measured field samples. AA and RB performed vulnerability curves measurements. AA and IR performed carbon isotope measurements. AA analyzed all data. AA, SC, and RDS co-wrote the manuscript with contributions from SD, HC, UH, and TK.

## Supporting Information

Table **S1**: Ψ_50_, Ψ_min_ and safety margins measured for the different species at the different sites in summer 2018.

Table **S2**: Ψ_50_, Ψ_min_ and safety margins measured for the different species at the different sites in summer 2019.

Table **S3**: Ψ_12_, Ψ_min_ and safety margins measured for the different species at the different sites in summer 2018.

Table **S4**: Ψ_12_, Ψ_min_ and safety margins measured for the different species at the different sites in summer 2019.

Table **S5**: Ψ_88_, Ψ_min_ and safety margins measured for the different species at the different sites in summer 2018.

Table **S6**: Ψ_88_, Ψ_min_ and safety margins measured for the different species at the different sites in summer 2019.

Table **S7**: Ψ_PD_ ANOVA analysis between sites per species per measurement date.

Table **S8**: Parameters for the linear regression of Ψ_MD_ vs. Ψ_PD_ for different species at different sites.

Table **S9**: Parameters for the linear regression of Ψ_MD_ vs. Ψ_PD_ for different species.

Table **S10**: Parameters correspond to Ψ_MD_ vs. П_0_, in all species at all sites.

Table **S11**: Ψ_PD_ ANOVA analysis between species per site per measurement date.

Table **S12**: Ψ_MD_ ANOVA analysis between species per site per measurement date.

Table **S13**: δ^13^C ANOVA analysis between species per site per measurement date.

Table **S14**: Summary of Covariance analysis testing influence of Ψ_MD_, site and Ψ_MD_ X site on Ψ_S_

Table **S15**: Summary of Covariance analysis testing influence of Ψ_MD_, site, and Ψ_MD_ X site, on П_0_

Table **S16**: Summary of Covariance analysis testing influence of Ψ_MD_, species and their interaction on П_0_

Table **S17**: Summary of Covariance analysis testing influence of Ψ_MD_, species and their interaction on Ψ_S_.

Table **S18**: Parameters correspond to Ψ_PD_ slopes analysis in all species at all sites.

Table **S19**: Measured and normalized parameters correspond to Fig. 7.

## Notes

### Competing Interest Statement

The authors have declared no competing interest.

## References

Ackerly DD. 2004. Adaptation, niche conservatism, and convergence: Comparative studies of leaf evolution in the California Chaparral. Ecological Monographs 163: 654–671.

Adams HD, Zeppel MJB, Anderegg WRL, Hartmann H, Landhäusser SM, Tissue DT, Huxman TE, Hudson PJ, Franz TE, Allen CD, et al. 2017. A multi-species synthesis of physiological mechanisms in drought-induced tree mortality. Nature Ecology and Evolution 1: 1285–1291.

Álvarez S, Rodríguez P, Broetto F, Sánchez-Blanco MJ. 2018. Long term responses and adaptive strategies of Pistacia lentiscus under moderate and severe deficit irrigation and salinity: Osmotic and elastic adjustment, growth, ion uptake and photosynthetic activity. Agricultural Water Management 202: 253–262.

Anderegg WRL. 2015. Spatial and temporal variation in plant hydraulic traits and their relevance for climate change impacts on vegetation. New Phytologist 205: 1008– 1014.

Anderegg WRL, Klein T, Bartlett M, Sack L, Pellegrini AFA, Choat B, Jansen S. 2016. Meta-analysis reveals that hydraulic traits explain cross-species patterns of drought-induced tree mortality across the globe. Proceedings of the National Academy of Sciences 113: 5024–5029.

Aranda I, Cadahía E, Fernández De Simón B. 2020. Specific leaf metabolic changes that underlie adjustment of osmotic potential in response to drought by four Quercus species. Tree Physiology 41: 728–743.

Bartlett MK, Klein T, Jansen S, Choat B, Sack L. 2016. The correlations and sequence of plant stomatal, hydraulic, and wilting responses to drought. Proceedings of the National Academy of Sciences 113: 13098-13103

Bartlett MK, Scoffoni C, Sack L. 2012. The determinants of leaf turgor loss point and prediction of drought tolerance of species and biomes: a global meta-analysis. Ecology Letters 15: 393–405.

Benito Garzón M, González Muñoz N, Wigneron J-P, Moisy C, Fernández-Manjarrés J, Delzon S. 2018. The legacy of water deficit on populations having experienced negative hydraulic safety margin. Global Ecology and Biogeography 27: 346–356.

Bittencourt PRL, Oliveira RS, da Costa ACL, Giles AL, Coughlin I, Costa PB, Bartholomew DC, Ferreira LV, Vasconcelos SS, Barros FV, et al. 2020. Amazonia trees have limited capacity to acclimate plant hydraulic properties in response to long-term drought. Global Change Biology 26: 3569–3584.

Canadell, J., R. B. Jackson, J. R. Ehleringer, H. A. Mooney, O. E. Sala and E.-D. Schulze 1996. Maximum rooting depth of vegetation types at the global scale. Oecologia, 108: 583–595.

Cernusak LA, Ubierna N, Winter K, Holtum JAM, Marshall JD, Farquhar GD. 2013. Environmental and physiological determinants of carbon isotope discrimination in terrestrial plants. New Phytologist 200: 950–965.

Chirino E, Bellot J, Sánchez JR. 2011. Daily sap flow rate as an indicator of drought avoidance mechanisms in five Mediterranean perennial species in semi-arid southeastern Spain. Trees 25: 593–606.

Choat B, Sack L, Holbrook NM. 2007. Diversity of hydraulic traits in nine Cordia species growing in tropical forests with contrasting precipitation. New Phytol 175: 686-698.

Choat B, Jansen S, Brodribb TJ, Cochard H, Delzon S, Bhaskar R, Bucci SJ, Feild TS, Gleason SM, Hacke UG, et al. 2012. Global convergence in the vulnerability of forests to drought. Nature 491: 752–755.

Cochard H, Damour G, Bodet C, Tharwat I, Poirier M, Améglio, T. 2005. Evaluation of a new centrifuge technique for rapid generation of xylem vulnerability curves. Physiol. Plant. 124: 410–418.

Cramer W, Guiot J, Fader M, Garrabou J, Gattuso J-P, Iglesias A, Lange MA, Lionello P, Llasat MC, Paz S, et al. 2018. Climate change and interconnected risks to sustainable development in the Mediterranean. Nature Climate Change 8: 972–980.

Creek D, Lamarque LJ, Torres-Ruiz JM, Parise C, Burlett R, Tissue DT, Delzon S. 2020. Xylem embolism in leaves does not occur with open stomata: evidence from direct observations using the optical visualization technique. Journal of Experimental Botany 71: 1151–1159.

Crombie D, Tippett J, Hill T. 1988. Dawn water potential and root depth of trees and understorey species in Southwestern Australia. Australian Journal of Botany 36: 621–631.

Danin A, Plitmann U. 1987. Revision of the plant geographical territories of Israel and Sinai. Plant Systematics and Evolution 156: 43–53.

Dawson TE, Mambelli S, Plamboeck AH, Templer PH, Tu KP. 2002. Stable Isotopes in Plant Ecology. Annual Review of Ecology and Systematics 33: 507–559.

Deligö A, Bayar E. 2018. Drought stress responses of seedlings of two oak species (Quercus cerris and Quercus robur). Turkish Journal of Agriculture and Forestry 42: 114–123.

Delzon S. 2015. New insight into leaf drought tolerance. Functional Ecology 29: 1247–1249.

Delzon S, Douthe C, Sala A, Cochard H. 2010. Mechanism of water-stress induced cavitation in conifers: Bordered pit structure and function support the hypothesis of seal capillary-seeding. Plant, Cell and Environment 33: 2101–2111.

Farquhar GD, Ehleringer JR, Hubick KT. 1989. Carbon isotope discrimination and photosynthesis. Annu Rev Plant Physiol Plant Mol Biol 40: 503-537.

Feng X, Ackerly DD, Dawson TE, Manzoni S, McLaughlin B, Skelton RP, Vico G, Weitz AP, Thompson SE. 2019. Beyond isohydricity: The role of environmental variability in determining plant drought responses. Plant Cell and Environment 42: 1104–1111.

Fontes CG, Dawson TE, Jardine K, McDowell N, Gimenez BO, Anderegg L, Negrón-Juárez R, Higuchi N, Fine PVA, Araújo AC, et al. 2018. Dry and hot: the hydraulic consequences of a climate change-type drought for Amazonian trees. Philosophical Transactions of the Royal Society B: Biological Sciences 373: 20180209.

Forner A, Valladares F, Aranda I. 2018a. Mediterranean trees coping with severe drought: Avoidance might not be safe. Environmental and Experimental Botany 155: 529–540.

Forner A, Valladares F, Bonal D, Granier A, Grossiord C, Aranda I. 2018b. Extreme droughts affecting Mediterranean tree species’ growth and water-use efficiency: the importance of timing. Tree Physiology 38: 1127–1137.

García de la Serrana R, Vilagrosa A, Alloza JA. 2015. Pine mortality in southeast Spain after an extreme dry and warm year: interactions among drought stress, carbohydrates and bark beetle attack. Trees 29: 1791–1804.

Gazol A, Sangüesa-Barreda G, Granda E, Camarero JJ. 2017. Tracking the impact of drought on functionally different woody plants in a Mediterranean scrubland ecosystem. Plant Ecology 218: 1009–1020.

González-Muñoz N, Sterck F, Torres-Ruiz JM, Petit G, Cochard H, von Arx G, Lintunen A, Caldeira MC, Capdeville G, Copini P, et al. 2018. Quantifying in situ phenotypic variability in the hydraulic properties of four tree species across their distribution range in Europe. PLoS ONE 13: e0196075.

Jakoby G, Rog I, Shtein I, Chashmonay I, Ben-Yosef D, Eshel A, Klein T. 2021. Tree Forensics: Modern DNA barcoding and traditional anatomy identify roots threatening an ancient necropolis. PLANTS, PEOPLE, PLANET 3: 211–219.

Johnson DM, Domec J-C, Carter Berry Z, Schwantes AM, McCulloh KA, Woodruff DR, Wayne Polley H, Wortemann R, Swenson JJ, Scott Mackay D, et al. 2018. Co-occurring woody species have diverse hydraulic strategies and mortality rates during an extreme drought. Plant, Cell & Environment 41: 576–588.

Johnson DM, McCulloh KA, Meinzer FC, Woodruff DR, Eissenstat DM, Phillips N. 2011. Hydraulic patterns and safety margins, from stem to stomata, in three eastern US tree species. Tree Physiology 31: 659–668.

Kolb KJ, Sperry JS. 1999. Differences in drought adaptation between subspecies of sagebrush (Artemisia tridentata). Ecology 80: 2373–2384.

Li X, Blackman CJ, Choat B, Medlyn BE, Rymer PD, Tissue DT. 2019. Drought tolerance traits do not vary across sites differing in water availability in Banksia serrata (Proteaceae). Functional plant biology : FPB 46: 624–633.

Lo Gullo MA, Salleo S. 1988. Different strategies of drought resistance in three Mediterranean sclerophyllous trees growing in the same environmental conditions. New Phytologist 108: 267–276.

Lobo A, Torres-Ruiz JM, Burlett R, Lemaire C, Parise C, Francioni C, Truffaut L, Tomášková I, Hansen JK, Kjær ED, et al. 2018. Assessing inter-and intraspecific variability of xylem vulnerability to embolism in oaks. Forest Ecology and Management 424: 53–61

Guo JS, Hultine KR, Koch GW, Kropp H, Ogle K. 2020. Temporal shifts in iso/anisohydry revealed from daily observations of plant water potential in a dominant desert shrub. New Phytologist 225: 713–726.

Jacobsen AL, Pratt RB, Davis SD, Tobin MF. 2014. Geographic and seasonal variation in Chaparral vulnerability to cavitation. Madroño 61: 317–327.

Klein T, Shpringer I, Fikler B, Elbaz G, Cohen S, Yakir D. 2013. Relationships between stomatal regulation, water-use, and water-use efficiency of two coexisting key Mediterranean tree species. Forest Ecology and Management 302: 34–42.

Lamy J-B, Delzon S, Bouche PS, Alia R, Vendramin GG, Cochard H, Plomion C. 2014. Limited genetic variability and phenotypic plasticity detected for cavitation resistance in a Mediterranean pine. New Phytologist 201(3): 874–886.

Larter M, Brodribb TJ, Pfautsch S, Burlett R, Cochard H, Delzon S. 2015. Extreme aridity pushes trees to their physical limits. Plant Physiology 168: 804–807.

Larter M, Pfautsch S, Domec J-C, Trueba S, Nagalingum N, Delzon S. 2017. Aridity drove the evolution of extreme embolism resistance and the radiation of conifer genus Callitris. New Phytologist 215: 97–112.

Liu C, Liu Y, Guo K, Fan D, Li G, Zheng Y, Yu L, Yang R. 2011. Effect of drought on pigments, osmotic adjustment and antioxidant enzymes in six woody plant species in karst habitats of southwestern China. Environmental and Experimental Botany 71: 174–183.

Maherali H, Pockman WT, Jackson RB. 2004. Adaptive variation in the vulnerability of woody plants to xylem cavitation. Ecology 85: 2184–2199.

Martin-StPaul N, Delzon S, Cochard H. 2017. Plant resistance to drought depends on timely stomatal closure (H Maherali, Ed.). Ecology Letters 20: 1437–1447.

Martínez-Vilalta J, Cochard H, Mencuccini M, Sterck F, Herrero A, Korhonen JFJ, Llorens P, Nikinmaa E, Nolè A, Poyatos R, et al. 2009. Hydraulic adjustment of Scots pine across Europe. New Phytologist 184: 353–364.

Meinzer FC, Johnson DM, Lachenbruch B, McCulloh KA, Woodruff DR. 2009. Xylem hydraulic safety margins in woody plants: Coordination of stomatal control of xylem tension with hydraulic capacitance. Functional Ecology 23: 922–930.

Meinzer FC, Woodruff DR, Marias DE, Smith DD, McCulloh KA, Howard AR, Magedman AL. 2016. Mapping ‘hydroscapes’ along the iso-to anisohydric continuum of stomatal regulation of plant water status. Ecology letters 19: 1343–1352.

Mencuccini M, Minunno F, Salmon Y, Martínez-Vilalta J, Hölttä T. 2015. Coordination of physiological traits involved in drought-induced mortality of woody plants. New Phytologist 208: 396–409.

Moreno-Gutiérrez C, Dawson TE, Nicolás E, Querejeta JI. 2012. Isotopes reveal contrasting water use strategies among coexisting plant species in a Mediterranean ecosystem. New Phytologist 196: 489–496.

Myers N, Mittermeler RA, Mittermeler CG, Da Fonseca GAB, Kent J. 2000. Biodiversity hotspots for conservation priorities. Nature 403: 853–858.

Nardini A, Casolo V, Dal Borgo A, Savi T, Stenni B, Bertoncin P, Zini L, McDowell NG. 2016. Rooting depth, water relations and non-structural carbohydrate dynamics in three woody angiosperms differentially affected by an extreme summer drought. Plant, Cell & Environment 39: 618–627.

Nemera DB, Bar-Tal A, Levy GJ, Lukyanov V, Tarchitzky J, Paudel I, Cohen S. 2020. Mitigating negative effects of long-term treated wastewater application via soil and irrigation manipulations: Sap flow and water relations of avocado trees (Persea americana Mill.). Agricultural Water Management 237: 106178.

Palacio S, Montserrat-Martí G, Ferrio JP. 2017. Water use segregation among plants with contrasting root depth and distribution along gypsum hills. Journal of Vegetation Science 28: 1107–1117.

Pammenter NW, Van der Willigen C. 1998. A mathematical and statistical analysis of the curves illustrating vulnerability of xylem to cavitation. Tree Physiology 18: 589–593.

Paudel I, Gerbi H, Wagner Y, Zisovich A, Sapir G, Brumfeld V, Klein T. 2019a. Drought tolerance of wild versus cultivated tree species of almond and plum in the field. Tree Physiology 40: 454–466.

Paudel I, Gerbi H, Zisovich A, Sapir G, Ben-Dor S, Brumfeld V, Klein T. 2019b. Drought tolerance mechanisms and aquaporin expression of wild vs. cultivated pear tree species in the field. Environmental and Experimental Botany 167: 103832.

Pratt RB, Jacobsen AL, Golgotiu KA, Sperry JS, Ewers FW, Davis SD. 2007. Life history type and water stress tolerance in nine california chaparral species (Rhamnaceae). Ecological Monographs 77: 239–253.

Pratt RB, Jacobsen AL, Jacobs SM, Esler KJ. 2012. Xylem transport safety and efficiency differ among Fynbos shrub life history types and between two sites differing in mean rainfall. International Journal of Plant Sciences 173: 474–483.

Rog I, Tague C, Jakoby G, Megidish S, Yaakobi A, Wagner Y, Klein T. 2021. Interspecific soil water partitioning as a driver of increased productivity in a diverse mixed Mediterranean forest. Journal of Geophysical Research: Biogeosciences 126: e2021JG006382

Redtfeldt RA, Davis SD. 1996. Physiological and morphological evidence of niche segregation between two co-occurring species of Adenostoma in California Chaparral. Écoscience 3: 290–296.

Rumman R, Atkin OK, Bloomfield KJ, Eamus D. 2018. Variation in bulk-leaf ^13^C discrimination, leaf traits and water-use efficiency–trait relationships along a continental-scale climate gradient in Australia. Global Change Biology 24: 1186–1200.

Serrano L, Peñuelas J, Romà Robert O, Serrano SL, Peñuelas J, Ogaya R, Savé R. 2005. Tissue-water relations of two co-occurring evergreen Mediterranean species in response to seasonal and experimental drought conditions. J Plant Res 118: 263– 269.

Skelton RP, Dawson TE, Thompson SE, Shen Y, Weitz AP, Ackerly D. 2018. Low vulnerability to xylem embolism in leaves and stems of North American oaks. Plant Physiology 177: 1066–1077.

Sofo A, Manfreda S, Fiorentino M, Dichio B, Xiloyannis C. 2008. Hydrology and Earth System Sciences The olive tree: a paradigm for drought tolerance in Mediterranean climates. Hydrolology & Earth System Sciences 12: 293–301.

Sparks JP, Black RA. 1999. Regulation of water loss in populations of Populus trichocarpa: the role of stomatal control in preventing xylem cavitation. Tree Physiology 19: 453–459.

Spinoni J, Vogt J V, Naumann G, Barbosa P, Dosio A. 2018. Will drought events become more frequent and severe in Europe? International Journal of Climatology 38: 1718–1736.

Stojnić S, Suchocka M, Benito-Garzón M, Torres-Ruiz JM, Cochard H, Bolte A, Cocozza C, Cvjetkovic B, de Luis M, Martinez-Vilalta J, et al. 2017. Variation in xylem vulnerability to embolism in European beech from geographically marginal populations. Tree Physiology 38: 173–185.

Tielbörger K, Bilton MC, Metz J, Kigel J, Holzapfel C, Lebrija-Trejos E, Konsens I, Parag HA, Sternberg M. 2014. Middle-Eastern plant communities tolerate 9 years of drought in a multi-site climate manipulation experiment. Nat Commun 5: 5102.

Toumi L, Lumaret R. 2010. Genetic variation and evolutionary history of holly oak: a circum-Mediterranean species-complex [Quercus coccifera L./Q. calliprinos (Webb) Holmboe, Fagaceae]. Plant Systematics and Evolution 290: 159–171.

Trumbore S, Brando P, Hartmann H. 2015. Forest health and global change. Science 349: 814–818.

Tyree MT, Sperry JS. 1989. Vulnerability of xylem to cavitation and embolism. Annual Review of Plant Physiology and Plant Molecular Biology 40: 19–36.

Tyree MT, Zimmermann MH. 1983. Xylem structure and the ascent of sap. NewYork, NY: Springer.

Väänänen PJ, Osem Y, Cohen S, Grünzweig JM. 2019. Differential drought resistance strategies of co-existing woodland species enduring the long rainless Eastern Mediterranean summer. Tree Physiology 4: 305–320.

Volaire F. 2018. A unified framework of plant adaptive strategies to drought: Crossing scales and disciplines. Global Change Biology 24: 2929–2938.

Wortemann R, Herbette S, Barigah TS, Fumanal B, Alia R, Ducousso A, Gomory D, Roeckel-Drevet P, Cochard H. 2011. Genotypic variability and phenotypic plasticity of cavitation resistance in Fagus sylvatica L. across Europe. Tree Physiology 31: 1175–1182.

Xu C, McDowell NG, Fisher RA, Wei L, Sevanto S, Christoffersen BO, Weng E, Middleton RS. 2019. Increasing impacts of extreme droughts on vegetation productivity under climate change. Nature Climate Change 9: 948–953.

Ziegler C, Coste S, Stahl C, Delzon S, Levionnois S, Cazal J, Cochard H, Esquivel-Muelbert A, Goret J-Y, Heuret P, et al. 2019. Large hydraulic safety margins protect Neotropical canopy rainforest tree species against hydraulic failure during drought. Annals of Forest Science 76: 115.

