## Supplemental Figure 1 for "Combined drought resistance strategies and the hydraulic limit in co-existing Mediterranean woody species"

### Slide 1
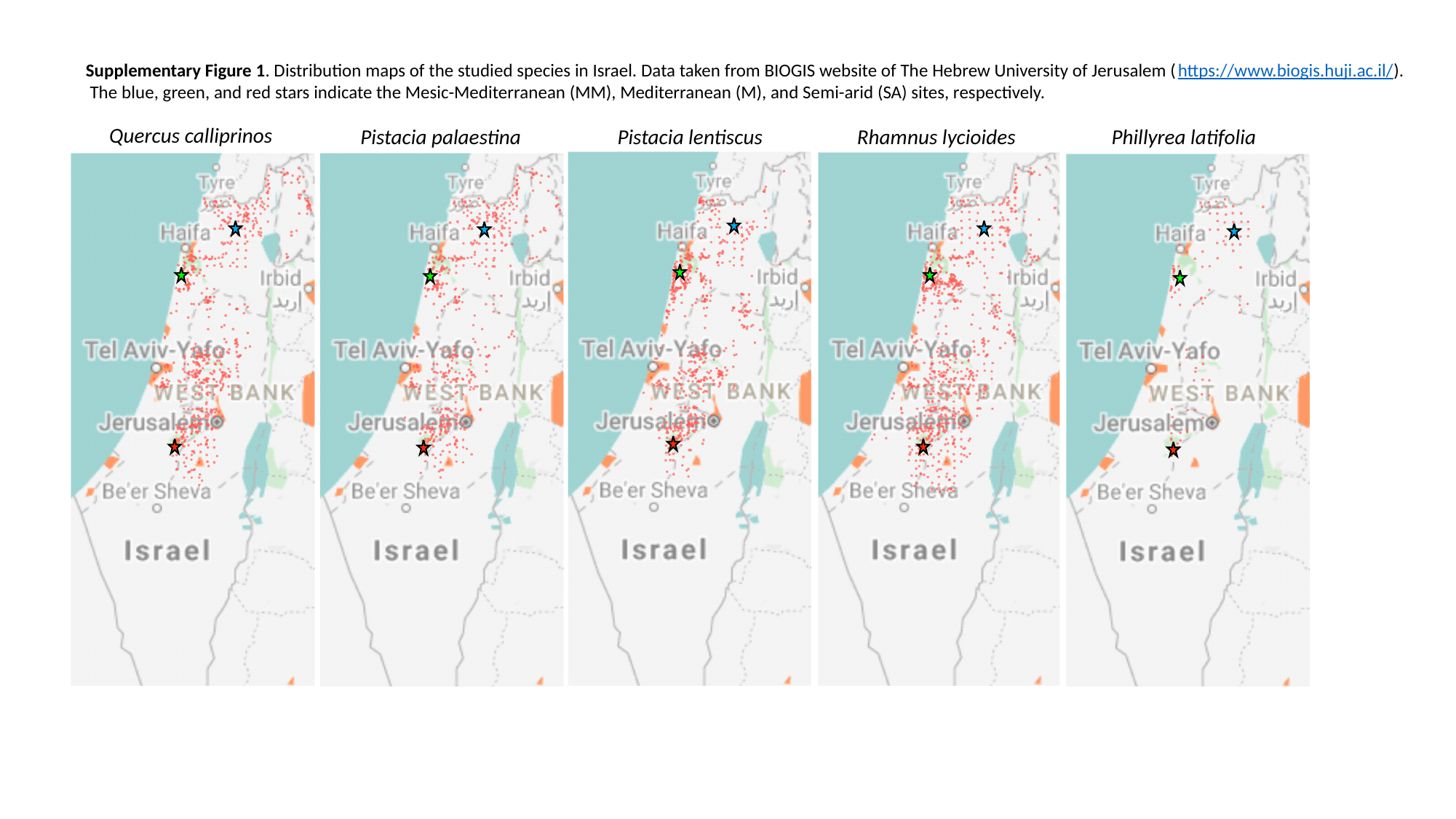

Supplementary Figure 1. Distribution maps of the studied species in Israel. Data taken from BIOGIS website of The Hebrew University of Jerusalem (https://www.biogis.huji.ac.il/).
 The blue, green, and red stars indicate the Mesic-Mediterranean (MM), Mediterranean (M), and Semi-arid (SA) sites, respectively.
Quercus calliprinos
Pistacia palaestina
Pistacia lentiscus
Rhamnus lycioides
Phillyrea latifolia
